# Evidence of structural rearrangements and variability in pESI(like) megaplasmids of *S*.Infantis by resolving the pESI-like ESBL-producing variants

**DOI:** 10.1101/2021.09.06.459061

**Authors:** Patricia Alba, Virginia Carfora, Fabiola Feltrin, Manuela Iurescia, Elena Lavinia Diaconu, Gessica Cordaro, Elena Dell’Aira, Ilaria Marani, Angelo Giacomi, Alessia Franco, Antonio Battisti

## Abstract

The increasing prevalence of pESI(like)-positive, multidrug-resistant (MDR) *S*. Infantis in Europe is a cause of major concern. As previously demonstrated, the pESI(like) megaplasmid is not only a carrier of antimicrobial resistant (AMR) genes (at least *tet, dfr* and *sul* genes), but also harbours several virulence and fitness genes, and toxin/antitoxin systems that enhance its persistence in the *S*. Infantis host.

In this study, five prototype pESI(like) plasmids, of either CTX-M-1 or CTX-M-65 ESBL producing strains, were long-read sequenced using Oxford Nanopore Technology (ONT) and their complete sequences were resolved. Comparison of the structure and gene content of the five sequenced plasmids, and further comparison with previously published pESI(like) sequences, indicated that although the sequence of such pESI(like) “mosaic” plasmids remains almost identical, their structures appear different and composed of regions inserted or transposed after different events.

The results obtained in this study are essential to better understand the plasticity and the evolution of the pESI(like) megaplasmid, and therefore to better address risk management options and policy decisions to fight against AMR and MDR in *Salmonella* and other food-borne pathogens.

## Introduction

*Salmonella enterica* subsp *enterica* serovar Infantis (*S*. Infantis) represents the fourth most commonly reported serovar in domestically acquired and travel-associated human infections and one of the top five serovars detected in food and food-producing animals in Europe, mostly related to broiler sources (93.1%) (EFSA and ECDC, 2021 a). Notably, *S*. Infantis accounted for most of the multidrug resistant (MDR) *Salmonella* spp. recovered from broilers and their derived carcases (79% and 75.3% respectively) (EFSA and ECDC, 2021 b). In Italy, MDR *S*. Infantis has increasingly been reported over the last decade from both food-producing animals and humans. This clone harbours a megaplasmid termed pESI-like (Franco et al., 2015; Alba et al., 2020), similar to the pESI (standing for plasmid of emerging *S*. Infantis) megaplasmid described in Israel (Aviv et al., 2014). Additionally, this pESI-like plasmid carried the ESBL gene *bla*_CTX-M-1_ (mediating extended-spectrum cephalosporin resistance, ESC-R), as well as resistance genes towards tetracyclines, trimethoprim, sulphonamides and aminoglycosides, heavy metals (*mer*A) and disinfectants (qacEΔ): Since then, it has been often detected in S. Infantis isolates from animal and human sources, in Italy and other European countries (Franco et al., 2015; Carfora et al., 2018; Alba et al., 2020). pESI-like plasmid is a conjugative megaplasmid acting as a parasite, as it harbours not only several AMR genes but also different toxin/antitoxin (T/AT) systems that have a key role in the permanence of the plasmid in the host and in the S. Infantis population (Franco et al., 2015; Alba et al., 2020). Therefore, the presence of pESI-like in *S*. Infantis represents a major concern as it often provides to zoonotic bacterial hosts additional traits of virulence, fitness and antimicrobial resistance (AMR), enhancing the rising prevalence of *S*. Infantis in broiler flocks and in human infections in Europe (EFSA and ECDC, 2021 b).

Importantly, another S. Infantis clone harbouring the pESI-like megaplasmid carrying the ESBL gene *bla*_CTX-M-65_, has been described in food-producing animals, and food thereof in the United States (Tate et al., 2017), where domestically acquired cases began in 2014 (Brown et al., 2018), and in few European human clinical cases mainly associated with a travel history to Asia and South America (Franco et al., 2015; EFSA and ECDC, 2021 b). In this regard, so far pESI-like *bla*_CTX-M-65_-positive *S*. Infantis appeared not to be associated with the European *S*. Infantis animal primary productions, and to date there is no evidence that it has entered the food-producing animal industry in the European Union (Alba et al., 2020).

Nevertheless, both “variants” of pESI-like included integrons associated to different AMR genes. Indeed, the presence of these genetic elements has recently demonstrated to accelerate the evolution from susceptible to resistant microorganisms (Souque, et al., 2021).

The improvement of third generation, high throughput sequencing (HTS) technology, or long-reads sequencing technology, has been proven useful for the study of the genetic structure of mosaic plasmids as pESI(like), because it overcomes the issues related with the assembly of regions containing insertion sequences (ISS) (Diaconu et al., 2020). Moreover, the use of a combined approach (e.g. Illumina-Oxford Nanopore Technologies, ONT, hybrid assembly) has been demonstrated to improve the accuracy and complete resolution of plasmid structures (George et al., 2017). Only in recent studies, few pESI-like plasmids have been complete sequenced (Cohen et al., 2020; Garcia-Soto et al., 2020, Kürekci et al., 2021; Tyson et al., 2020) but none of them belonged to the *bla*_CTX-M-1_ variant, that has colonised the European *S*. Infantis population (Alba et al., 2020).

The aims of this study were: a) to fully reconstruct and resolve four representative pESI-like-*bla*_CTX-M-1_ positive plasmids from animal and food sources collected at the IZSLT in 2011-2016 and one pESI-like - *bla*_CTX-M-65_ positive plasmid identified in 2014 from an European (Italian) human patient with a travel history to America; b) to gain insight into differences and similarities of the studied plasmids and other complete publicly available pESI-like plasmids by comparing the structure, the coding regions and their nucleotide sequences using a hybrid assembly (Illumina Oxford Nanopore Technologies, ONT) approach, with a particular focus on insertions, deletions and rearrangements

## Material and Methods

### S. Infantis isolates

Five *S*. Infantis isolates, collected by the National Reference Laboratory for Antimicrobial Resistance (NRL-AR), Istituto Zooprofilattico Sperimentale del Lazio e della Toscana (IZSLT) ‘M. Aleandri’ in 2016-2017 and already phenotypically and genetically characterized also by Illumina short-reads sequencing (Franco et al., 2015, Carfora et al., 2018; Alba et al., 2020), were selected to be further sequenced with long read approach (ONT). These five isolates, whose metadata are summarized in Table1, were selected for the following reasons:

- three isolates (IDs: 11024547-4, 12037823/11 and 13065790-183) were the oldest available in the collection having pESI-like encoding *bla*_CTX-M-1_ from meat from pigs and broilers.
- one isolate (ID: 14026835) harbouring bla_CTX-M-65_ was very different from the main pESI-like plasmid circulating in Italy, according to all the previous studies (Franco et al., 2015, Alba et al., 2020)
- one isolate (ID: 16092401-41) was characterized by the co-presence in the host cell (*S*. Infantis) of *bla*_CTX-M-1_-positive-pESI-like together with an IncX1 plasmid carrying *mcr*-1 colistin-resistance gene.

**Table 1.**
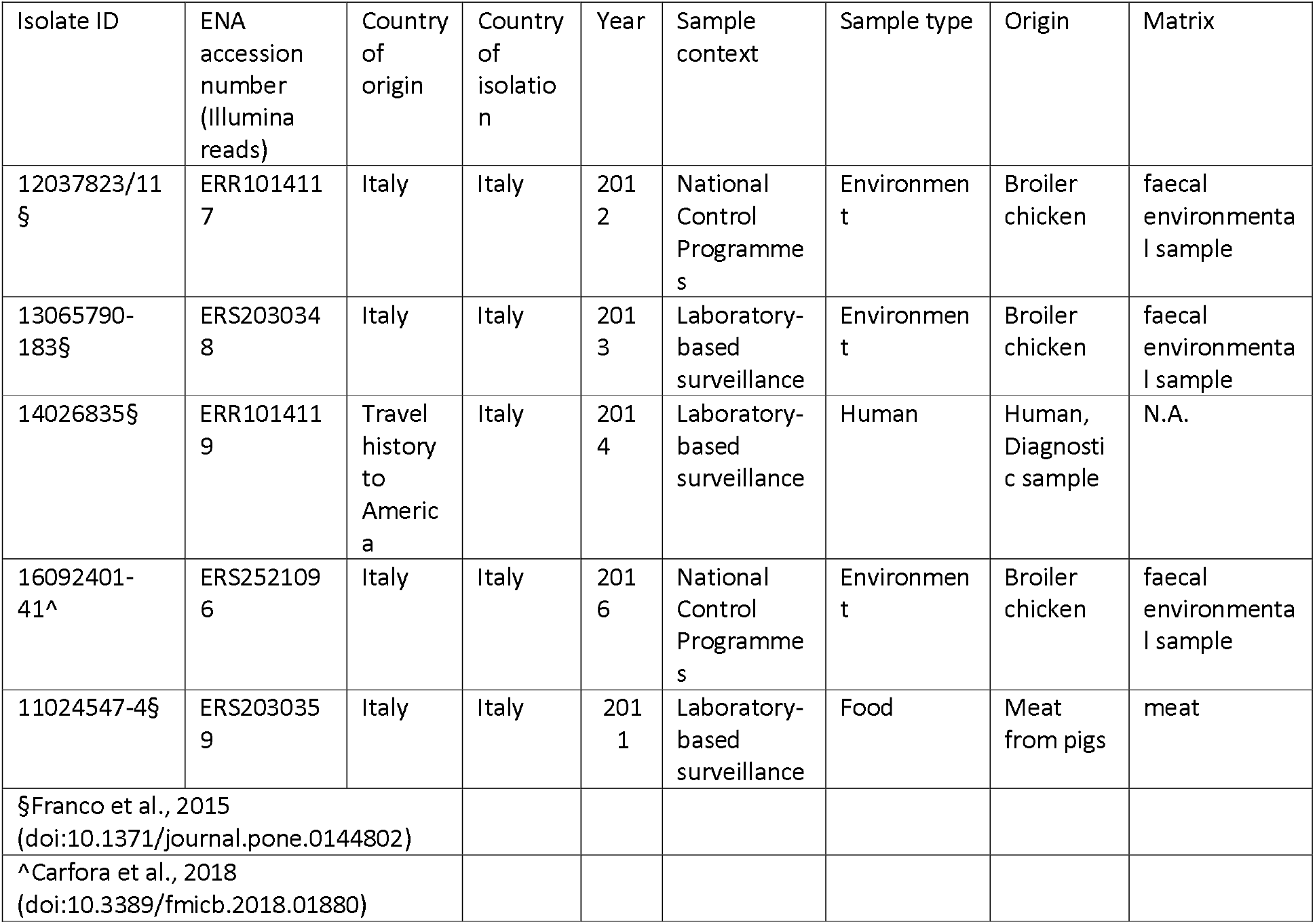
General isolate information

| Isolate ID | ENA accession number (Illumina reads) | Country of origin | Country of isolation | Year | Sample context | Sample type | Origin | Matrix |
| --- | --- | --- | --- | --- | --- | --- | --- | --- |
| 12037823/11§ | ERR1014117 | Italy | Italy | 2012 | National Control Programmes | Environment | Broiler chicken | faecal environmental sample |
| 13065790-183§ | ERS2030348 | Italy | Italy | 2013 | Laboratory-based surveillance | Environment | Broiler chicken | faecal environmental sample |
| 14026835§ | ERR1014119 | Travel history to America | Italy | 2014 | Laboratory-based surveillance | Human | Human, Diagnostic sample | N.A. |
| 16092401-41^ | ERS2521096 | Italy | Italy | 2016 | National Control Programmes | Environment | Broiler chicken | faecal environmental sample |
| 11024547-4§ | ERS2030359 | Italy | Italy | 2011 | Laboratory-based surveillance | Food | Meat from pigs | meat |
| §Franco et al., 2015<br>(doi:10.1371/journal.pone.0144802) |  |  |  |  |  |  |  |  |
| ^Carfora et al., 2018<br>(doi:10.3389/fmicb.2018.01880) |  |  |  |  |  |  |  |  |

### Sequencing and bioinformatics analysis

Genomic DNA was extracted by using QIAamp DNA Mini Kit (Qiagen, Hilden, Germany) following the manufacturer’s protocol.

Libraries for Oxford Nanopore Technologies (ONT) were prepared with the rapid barcoding kit (SQK-RBK004) and sequenced using the nanopore-based MinION device (Branton D. et al, 2008). A hybrid (lllumina-ONT) assembly was carried out using the Unicycler pipeline (Wick et al., 2017) with the default parameters.

Annotation was carried out using RAST (Azik et al., 2008) and curated using Prokka (Seeman et al., 2014), ResFinder (Bortolaia et al., 2020), PlasmidFinder (Carattoli et al., 2014), and ISFinder (Siguier et al., 2008) tools. Blast (Zhang et al., 2000) on line (nr. database) was also used for specific regions.

Graphical representation of the general structures and genetic regions was performed by using GView (https://www.gview.ca/; Petkau et al., 2010) and Artemis Comparison Tool (ACT) (Carver et al., 2005). The complete sequences of the plasmids have been deposited in the European Nucleotide Archive (ENA) at EMBL-EBI under accession number PRJEBXXXXX.

All plasmids were compared with the pESI-like sequence from 12037823, used as reference.

The complete five pESI-like plasmid sequences obtained were also compared with two complete pESI-like plasmid sequences previously obtained with long reads technologies and publicly available, as CP047882 (Cohen et al., 2020) and NZ_CP016409 (Tate et al., 2017).

## Results

### Common structural regions of pESI-like plasmids

Five selected pESI-like plasmids have been complete sequenced and resolved. All plasmids studied consisted in a mosaic structure: they contained *rep*B, that according to PlasmidFinder belonged to the IncFIB replicon type (1..1001 nt.). They also Included a non functional IncP origin of replication (176043..176934 nt.; Identity 100%; coverage 82%) and almost the complete sequence of the IncI plasmid (182475..263544 nt; Identity 90%; coverage 96.56%; AP005147.1), except the region enconding the IncI replication origin (Figure 1)

**Figure 1:**
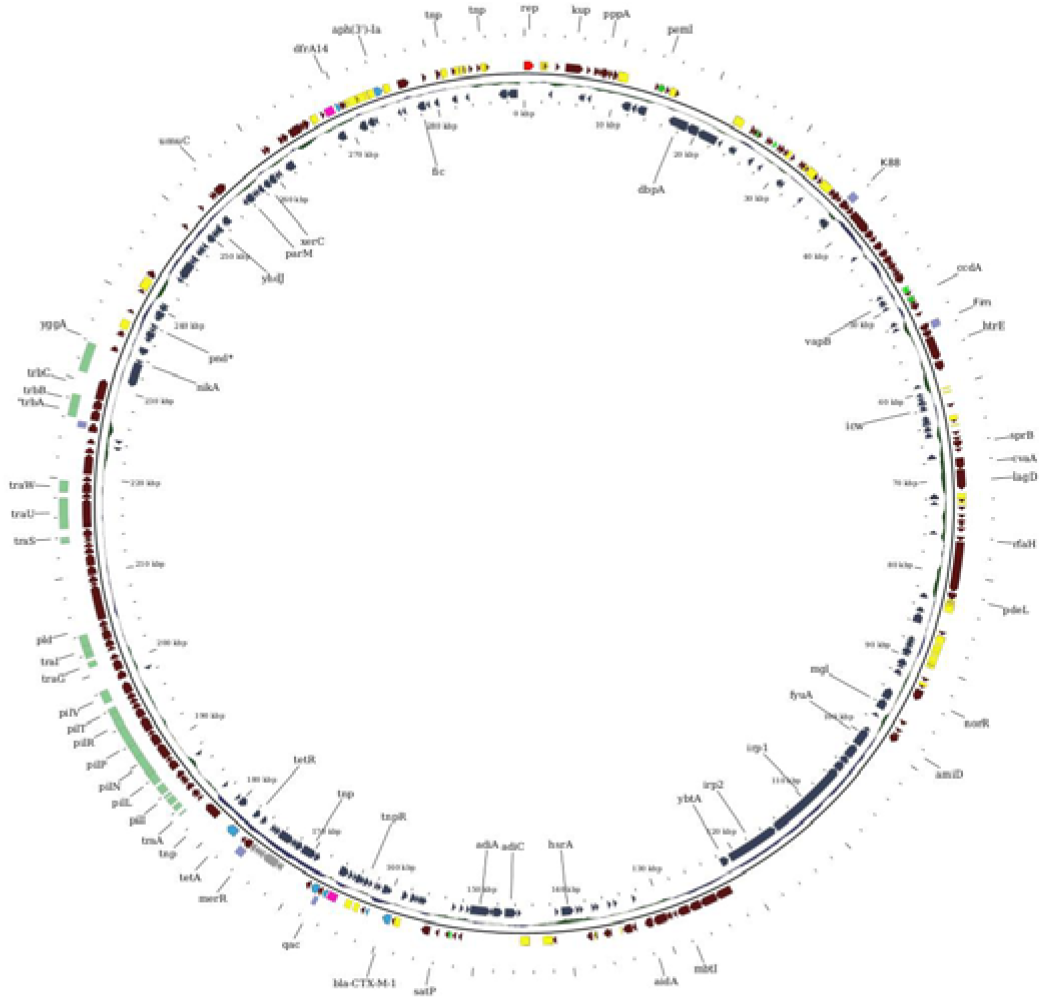
Graphical representation of the pESI-like plasmid complete sequence (ID 12037823/11). Block colours indicate the function of the genes: red: *rep*B gene; pink: class I integrons (IntI); yellow; mobile elements; blue: resistance genes; green: toxin/anti-toxin systems; grey: mer operon; light green: genes involve in conjugation.

The size of the pESI-like plasmid found in the isolate ID 12037823/11 from a broiler sample had a 289912 bp size and, according to the annotation contained 402 Coding sequences (CDS) and 50 of them were classified as mobile elements. The five pESI-like plasmids contained a different number of CDS, in particular 400 CDS for ID 11024547, 418 CDS for ID 13065790, 438 CDS for ID 14026835 and 404 CDS for ID 16092401.

The accurate annotation of pESI-like from isolate ID 12037823/11 revealed the presence of two integrons: one contained an integrase (Int)1 (165385..166398) associated to *bla*_CTX-M-1_, *dfr*1, qacΔE1 and *sul*1 genes (166555..168400) and one an Int2 (268153..269130) associated to *dfr*A14 (269452..269934). Moreover, besides the presence of the T/AT systems PemI/K (14531..15121) and CcdA/B (49897..50422) previously described, VapB/C (48630..49273) and HigA modules have been also found (152724..153032). Moreover, there was evidence of other T/AT systems relics, such as antitoxin DinJ (25860..26147) without the presence of the YafQ toxin, and a sequence 86% similar to the post-segregation killing protein PndC gene (237758..238192) of the plasmid pIFM3804 of *E. coli* (KF787110) without the corresponding *pnd*B (Figure 1)

This megaplasmid also harboured the yersiniabactin operon (99383..128414) with a percentage of identity of 97.4% with the yersiniabactin operon of *Yersinia pestis* (AF091251), the complete mercury resistance (mer) operon (172047..176009) and the ImpABC Operon (253221..255221) (Figure 1).

### Comparison of the plasmid structures

The two by two comparison of the genetic structure of the five plasmids, by using the ACT tool, indicated the presence of structural differences. As for the two isolates from broiler chicken samples of 2012 and 2013, the pESI-like plasmid of isolate ID 13065790 was slightly different and 8980 bp larger than that in the isolate ID 12037823. In particular, a 6000 bp long fragment (160303..166308 in pESI-like of isolate 13065790) bracketed with IS26 mobile elements resulted inserted. This insertion was composed by a duplication of the IS26 in pESI-like in the ID 13065790, surrounding a truncated additional Int1 (160348..166263) and the following genes: *aad*A1-like (160289..161079), *tnp* (161317..161892), *sul3* (162250..163041), short-chain dehydrogenase (163793..164656) and *mef*(B) (164802..166031). pESI-like from ID13065790 also included a second *aad*A1-like gene in the position 175255..176046 (corresponding to position 167106 of pESI-like from 12037823).

As for pESI-like in ID 12037823 (289912 bp) and pESI-like in 11024547 (290187 bp), they were very similar but two differences were observed: a) a deletion and rearrangement in an autotransporter outer membrane beta-barrel domain-containing protein (76508..81841 of 12037823; 76508..81235 of 11024547) and b) a new *aad*A1-like gene (166554..167345 of 11024547) in the position 167114 of pESI-like 12037823.

The obtained sequences and structures of the pESI-like plasmids in isolates ID 12037823 and ID 16092401 indicated that both plasmids were identical (Figure 2).

**Figure 2:**
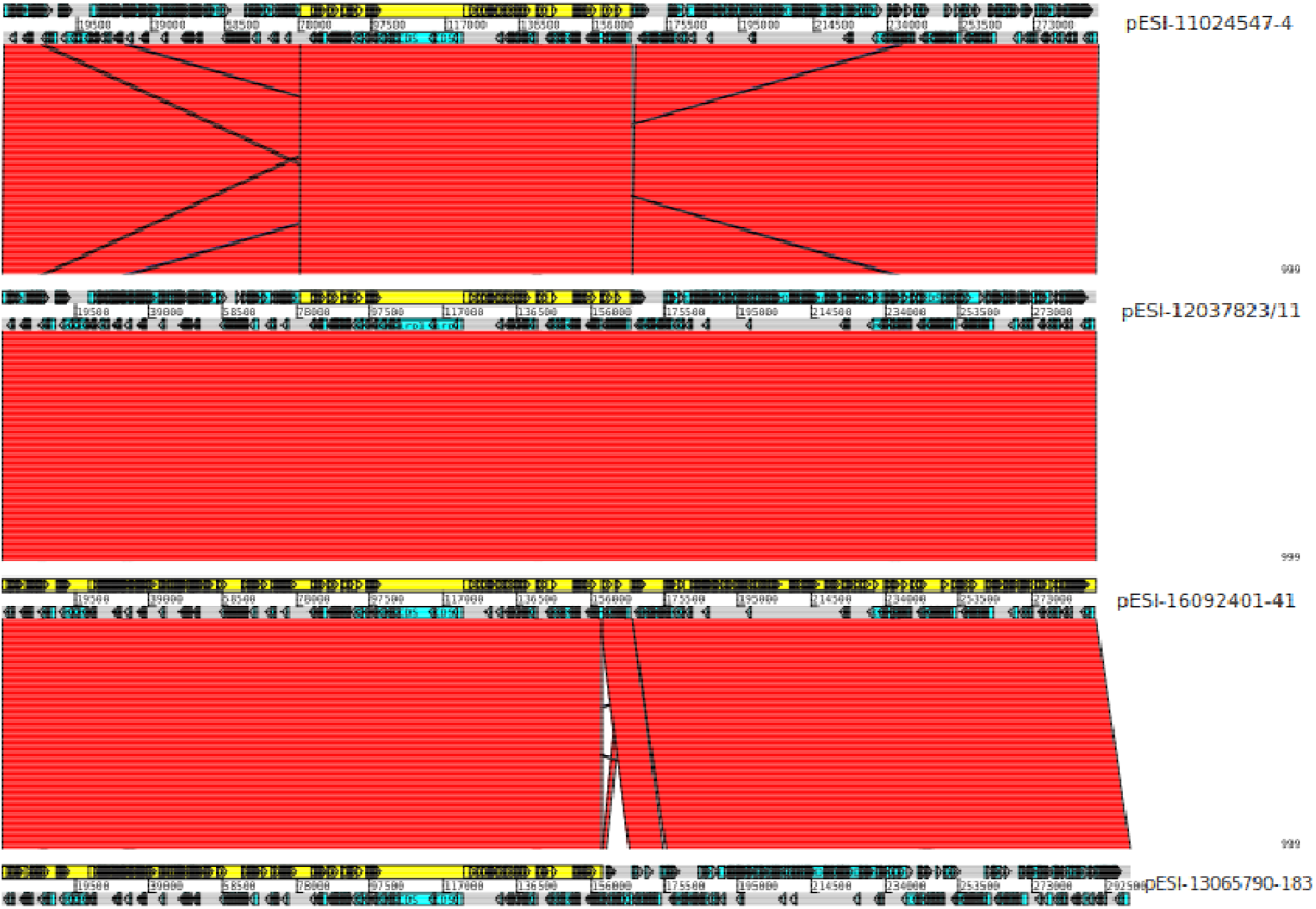
Comparison of four complete sequences of pESI-like plasmids harbouring *bla*_CTX-M-1_ from Italian isolates IDs 11024547-4, 12037823/11, 16092401-41 and 13065790-183, using ACT. Red Squares indicated 99% of similarity and the same sense of the sequence.

Differently, the *bla*_CTX-M-65_-positive pESI-like from ID 14026835 (323674 bp) presented the highest number of differences when compared with that in ID 12037823. Regarding the structure, we have found different situations: a) homologue regions in the same position and sense, as the replication region (1..23412; 274744..289902; containing *rep, parA* and *parB* (Figure 2).

b) homologue regions in different relative positions but in the same sense as the region that included the nickel transporter codon, two T/AT systems and the pESI(like) markers K88 and Fim (23404..54861 of pESI-like in ID 12037823 corresponds to 251799..283256 of pESI-like in ID 14026835);

c) homologue regions in different relative position and in the inverse sense as the region where *qac* and *sul* genes were located (167106..169132 of pESI-like in ID 12037823; 139947..137921 of pESI-like in ID 14026835) or the region containing the mer operon and the *tet*A gene (169129..197414 of pESI-like from 12037823; 135049..106264 of pESI-like from 14026835).

Importantly, the *bla*_CTX-M-65_-positive pESI-like in ID 14026835 also presented differences in the gene content. In particular, it had lost several regions containing *mphA* (160182..165326), *dfrA1* (166539..167105), or the plasmid stability gene parM (255122 262514) when compared with the “European” pESI-like in ID 12037823. On the other side, it had incorporated several regions that harboured the arsenic resistance genes (23412..30499), the Hok post-segregation killing protein (75492..77469) gene and a multidrug resistance region containing *aac*(3)-IVa, *aph*(4)-la, *floR*, and *fosA3* (293012..308516). Finally, in terms of resistance towards expanded-spectrum cephalosporin resistance, the most relevant difference we observed was the presence of *bla*_CTX-M-1_ in pESI-like in ID 12037823 and *bla*_CTX-M-65_ in pESI-like in ID 14026835, albeit situated in different plasmid locations, 159012..160181 and 250838..249707, respectively (Figure 2).

### Comparison of pESI-like from ID 12037823 and pESI from CP047882

Results from the comparison of pESI-like in ID 12037823 and pESI in CP047882 (Cohen et al., 2020), the first described pESI megaplasmid identified in Israel, indicated the presence of common regions intercalated with inserted sequences/transposons (Figure 3). One of this transposon was the region where *bla*_CTX-M-1_ was located (158111..165150) in the pESI-like in ID 12037823, inserted in the position 158169..158988, while pESI-like in CP047882 presented a ISS element. Other differences were the presence of *aad*A1 (160380..161225) in pESI-like in CP047882, in a region where pESI-like in ID 12037823 (166539..167105) harboured *dfr*, or the inserted regions where *aph*(3’)-la was located in pESI-like in ID 12037823 (273748..274871), that was absent in pESI-like in CP047882 (Figure 3)

**Figure 3:**
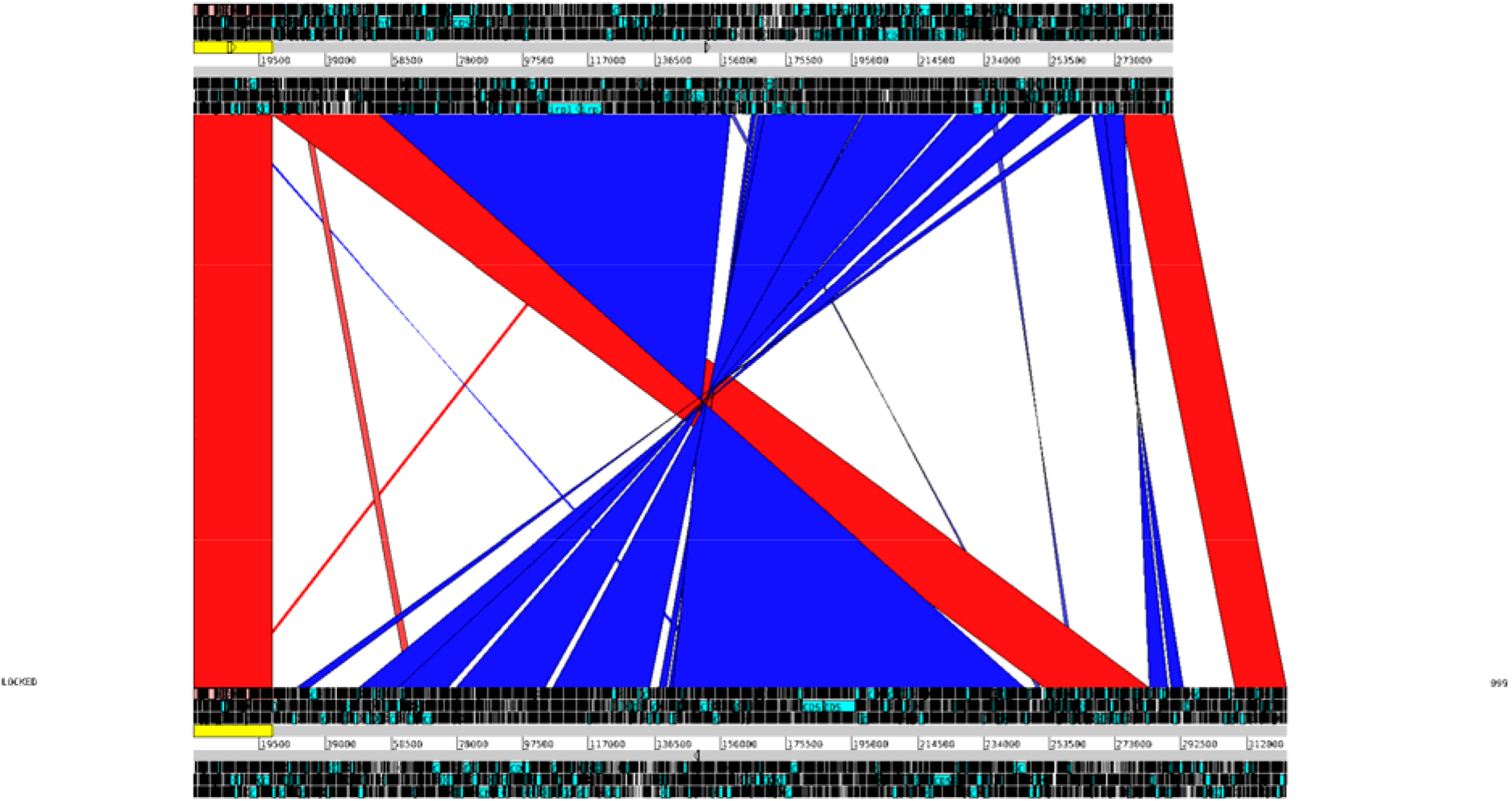
Comparison of pESI-like from ID 12037823 (CTX-M-1-positive, Italy) and pESI from CP047882 (CTX-M-1-negative, Israel). Red Squares indicated 99% of similarity and the same sense of the sequence. Blue squares indicated 99% of similarity but inverted sequences.

### Comparison of pESI-like from 12037823 and pESI-like from NZ_CP016409.1

The comparison of pESI-like in ID 12037823 and pESI-like in NZ_CP016409.1 harbouring *bla*_CTX-M-65_(Tate et al., 2017) revealed the insertion, deletion and rearrangements of regions containing important genes.

In the region 159013..165326 of pESI-like in ID 12037823, it was located the ESBL gene *bla*_CTX-M-1_ that was inserted in the corresponding position 159028 nt. of pESI-like in NZ_CP016409.1. The range 165327.. 166538 of pESI-like in ID 12037823 corresponded to 159027..160241 of pESI-like in NZ_CP016409.1, but in the first plasmid was located the gene *dfr* whereas in the second plasmid were located the genes *ant*3”-la and *aad*A.

A 27887 bp region (270536..298423) of pESI-like in NZ_CP016409.1 corresponded to a 134 bp region of pESI-like in ID 12037823. In this insertion sequence of the NZ_CP016409.1’s plasmid were located several resistance genes to antimicrobials and heavy metals including *ars*H, sulfite exporter encoding gene, *bla*_CTX-M-65_, *fos*A, *tet*R, *flo*R, *aph*(4)-la and *aac*(3)-lva.

### Identical sequence but different structure: comparison of pESI-like plasmids harbouring bla_CTX-M-65_

When comparing the sequences of pESI-like described by Tate et al. (NZ_CP016409) of 322518 bp and the pESI-like from isolate ID 14026835 of 323674 bp, both harbouring the <u>*bla*_CTX-M-65_</u> gene, we found that they shared a 99% of identity. The different size of both plasmids was due mainly to an additional IS15 transposon of 780 bp (251020..251799; IS6 family) in pESI-like from ID 14026835.

The comparison of plasmid structures indicated that the replication region of the plasmid, around 37400 bp, remained constant in both, sequence and structure (308516..23412 nt; NZ_CP016409 position). The rest of the plasmid, was almost identical (>99% identity), but the presence of a transposition of one region (251799..283256 region of pESI-like in NZ_CP016409 and 23404..54861 region of pESI-like in 14026835) and the inversion of three regions (246185..106264 region of pESI-like in NZ_CP016409 that correspond to 54854..194771 region of pESI-like in 14026835; 105200..23412 region of pESI-like in NZ_CP016409 that correspond to 195835..277623 region of pESI-like in 14026835; 309339..283251 region of pESI-like in NZ_CP016409 that correspond to 282097..308185 region of pESI-like in 14026835) resulted in two identical plasmid with inverted structures (Figure 4).

**Figure 4.**
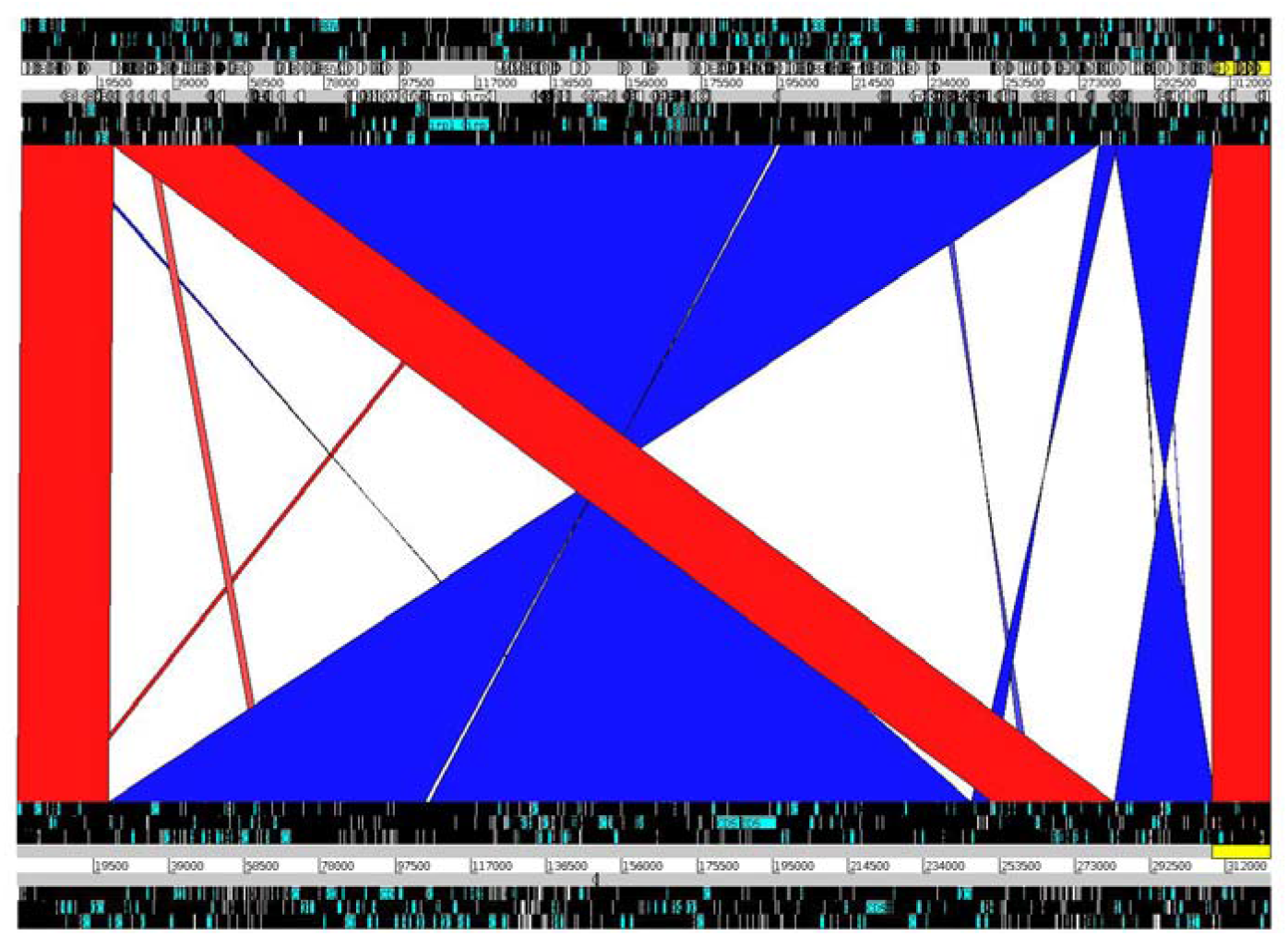
Comparison of pESI-like from NZ_CP016409 (CTX-M-65-positive, USA) and pESI-like from isolate ID 14026835 (CTX-M-65-positive, Italy, travel history to America). Red Squares indicated 99% of similarity and the same sense of the sequence. Blue squares indicated 99% of similarity but inverted sequences.

## Discussion

In the present study, the structures of five representative pESI-like megaplasmids from ESBL-producing *S*. Infantis were resolved using a hybrid (lllumina-ONT) sequencing approach and the resolved plasmidic structures were two by two compared. This study indicates that large plasmids could be subject to recombinations and rearrangements in their structure, most likely due to the selective pressure of the farm environment, still maintaining all the genetic determinants and the capability of acquiring additional genetic elements. However, also considering the main findings of this study, the general question of whether these megaplasmids have a different evolutionary origin or are the results of consecutive rearrangements events, remains unanswered.

We observed that the first pESI described, in Israel (Aviv et al., 2014, Cohen et al., 2020), was smaller than the pESI-like described in Italy. In particular, a large part of the Italian pESI-like plasmid population has acquired a transposon containing *bla*_CTX-M-1_, as previously described (Franco et al., 2015; Carfora et al., 2018; Alba et al., 2020). This transposon has been integrated close to a Class-I integron (IntI) together with *dfr*1, *qac*ΔE1 and *sul*1 resistance genes. Differently, the pESI-like described in the United States (Tate et al., 2017), when compared with the pESI from Israel, has integrated a larger region including *bla*_CTX-M-65_ together with other resistance genes towards antibiotics (e.g. *floR, fosB*) or heavy metals (*arsH*).

The megaplasmids classified as pESI-like could be described as a mosaic plasmid, the result of the fusion, rearrangements or transposon integration of different plasmid types. Most importantly, the elevate number of resistance genes (*bla*_CTX-M-1_, *dfr*1, *qac*ΔE1, *sul*1) that pESI-like has acquired in different moments and in different positions, have conferred to pESI-like particular MDR traits. Moreover, it harboured other plasmid-borne virulence factors as the yersiniabactin operon, the mer operon and the ImpABC operon which contains a functional UmuDC error-prone DNA repair system involved in the resistance to the UV irradiation (Runyen-Jnecky, et al., 1999).

Another important feature of this plasmid is the capability of preventing its cure by the host, through post-segregation killing activity of the different T/AT systems (PemI/K, CcdA/B, VapB/C, HigA) as previously described (Alba et al. 2020).

The detailed description of five pESI-like infecting S. Infantis isolated from different sources and years in Italy, highlights the plasticity of this plasmid to adapt itself and its host to different environmental conditions and selection pressure. Indeed, only two of the analysed plasmids, both from S. Infantis isolated from broiler sources but in different years (2012 and 2016), could be considered identical. Moreover, a high chromosomal similarity between this two isolates was observed in previous studies (Alba et al., 2020). Those observations do not suggest important microevolution changes of both core genome and pESI-like plasmid along the 4-year interval within the same production system environment (Italian broiler chicken industry). Noteworthy, however, the S. Infantis host of 2016 also presented and IncX plasmid harbouring the mcr-1 transferable gene (Carfora, et al., 2018), so that in this case the strain may have microevolved by the acquisition of a new plasmid following the selection pressure exerted during this time interval by the extensive use of colistin. As expected, the pESI-like from the bla-CTX-M-65 producing S. Infantis of human origin was the most divergent from the other four bla-CTX-M-1-positive pESI-like megaplasmids sequenced, due to the presence of additional resistance (floR, fosA, aph(4)-la) and virulence genes acquired by insertions, rearrangements or inversions. These findings are in agreement with its epidemiology and phylogeography associated with the Americas (Alba et al, 2020) and with “the modular nature” of pESI observed by Cohen et al., 2020. Interestingly, recent studies showed that pESI-like could also recombine with IncX plasmids (Kürekci et al., 2021), confirming its overall plasticity.

Overall, the results of this study represent further evidence of the importance to exploit long sequencing methods to study the plasmid sequences and structure variation, as they are helpful to avoid some of the challenges raised by the study of mosaic plasmids (Pesesky M., et al, 2019). pESI-like has been described in S. Infantis worldwide, e. g. in Israel (Aviv et al., 2014), Italy (Franco et al., 2015), United States (Tate et al., 2017), Switzerland (Hindermann et al., 2017), Japan (Yokoyama et al., 2015), Hungary (Szmolka et al., 2018), Brazil (dos Santos et al., 2021) and Turkey (Kürekci et al., 2021). With few exceptions, in these studies the characterization of the plasmids was done using short reads sequencing, and the results indicated a less diversity between them compared to what has been observed when the plasmids were fully resolved. Even in a wide European study including outgroup genomes from the Americas, using short reads (Alba, et al. 2020), the main variability was related to the presence of different bla_CTX-M_ encoding genes (bla_CTX-M-1_ vs bla_CTX-M-65_) or other marker genes (e. g. *fosB*).

Moreover, application of long reads sequencing is also useful to identify repeated genes, as the case of isolate ID 13065790, that presented a duplicated IntI and *aad*A1. Otherwise, as previously published (Alba et al., 2020), if different variants of the same gene are present, assembly of short reads can affect negatively the detection and identification of those genes.

Based on the results herein described and in other previous studies (Tate et al., 2017, Gymoese et al. 2019, Kürekci et al., 2021), we proposed two main hypothesis that could explain the diversity of the pESI-like plasmids analysed: i) the modification of one initial megaplasmid with different rearrangements and inversion or ii) a convergent plasmid evolution in different S. Infantis strains following the fusion of smaller plasmids previously present in the bacterial hosts, as already observed in *B. cereus* (Zheng et al 2013). These *B. cereus* plasmids have been further selected because of the evolutionary advantage in the prokaryotic cell, probably because positive selection has been reported to favour a reduced plasmid variability (Carrilero et al., 2021). The answer on whether these megaplasmids have different evolutionary origin or are the results of consecutive rearrangements events remains elusive, Still, our study provides important insights into the structure of these complex genetic elements, which are of input for phylogeny, phylogeography, genomic epidemiology and source attribution purposes.

However, further studies should be carried out to better understand the drivers and evolution mechanisms behind the emergence of such complex plasmids. These studies may include the investigation of possible compensatory evolution events in the chromosome that could positively or negatively affect the presence of these plasmids in case of variation in fitness of Salmonella strains in particular environmental conditions, as in antibiotic-free environments. In some cases, gene loss at chromosome level may also occur to improve fitness, when host-parasite interaction with the plasmid and its genes composition has become stable (Hall et al., 2020). Investigating the structure and origin of MDR-megaplasmids as pESI-like of *S*. Infantis and its parasitic relationships with the host, would help us to prevent in the future the emergence and spread of new megaplasmids harbouring virulence genes and AMR genes to Highest Priority Critically Important Antimicrobials (HPCIAs).

## Acknowledgements

The genomic work was conducted in the framework of the Full Force project, supported by funding from the European Union’s Horizon 2020 Research and Innovation programme under grant agreement No 773830: One Health European Joint Programme.

## Author contribution

Conceptualization: PA, VC, AF, AB; Wet Lab analysis: FF, GC, ED, IM, AG; Bioinformatic analysis: PA, ELD; Investigation: PA; Resources: AF, AB; Data curation: PA; Writing of original draft preparation: PA, VC; Writing of review and editing: all authors. Visualization: PA Supervision: AF, AB.

